# Submedius Thalamus Modulates Orbitofrontal Cortex Representations During Maternal Behavior in Mice

**DOI:** 10.1101/2025.09.18.677207

**Authors:** Gen-ichi Tasaka, Mitsue Hagihara, Haruna Kobayashi, Miho Kihara, Takaya Abe, Kazunari Miyamichi

**Affiliations:** Laboratory for Comparative Connectomics, RIKEN Center for Biosystems Dynamics Research, Kobe, Hyogo 650-0047, Japan; Japan Science and Technology Agency, PRESTO, Kawaguchi, Saitama 332-0012, Japan; Laboratory for Animal Resources and Genetic Engineering, RIKEN Center for Biosystems Dynamics Research, Kobe, Hyogo 650-0047, Japan

## Abstract

The orbitofrontal cortex (OFC) is central to cognitive and social functions, yet its presynaptic partners remain incompletely defined. In female mice, the OFC modulates infant-directed maternal caregiving behaviors essential for offspring survival in an experience-dependent manner. Here, we identify the submedius thalamus (SMT) as a major presynaptic partner of the OFC. Trans-synaptic tracing revealed intensive inputs from both the SMT and mediodorsal thalamus (MD) to OFC layer 5 excitatory neurons. A mouse line *Tnnt1-Cre* enabled selective targeting of these higher-order thalamic nuclei. Axonal tracing demonstrated complementary projection patterns of SMT and MD across prefrontal regions. Microendoscopic Ca^2+^ imaging demonstrated pup retrieval encoding in SMT and MD, but only SMT exhibited learning-related plasticity, characterized by enhanced anticipatory responses as acquired maternal behaviors. Projection-specific chemogenetic silencing demonstrated that only SMT modulates OFC activity during pup retrieval. These findings demonstrate SMT as a previously uncharacterized thalamic hub shaping cortical representations of maternal behavior.

**Highlights:**

- OFC^Rbp4^ neurons receive strong inputs from higher-order thalamic nuclei (SMT, MD).
- SMT and MD show complementary axonal projections across prefrontal cortical cortex.
- SMT ensembles encode pup retrieval with experience-dependent plasticity.
- SMT inputs selectively modulate pup retrieval representation in OFC^Rbp4^ neurons.

## INTRODUCTION

The orbitofrontal cortex (OFC), a subdivision of the prefrontal cortex (PFC), has attracted growing interest for its role in regulating cognitive functions and social behaviors. Conceptually, the OFC has been proposed to construct a cognitive map of task states, an abstract representation of currently relevant features that supports inference and learning.^1–4^ It also plays an important role in linking outcomes to preceding choices, enabling rapid reversal, devaluation, and flexible updating of stimulus-outcome contingencies.^5–10^ Recent studies demonstrate that individual OFC neurons encode a wide range of behaviorally relevant features, including sensory inputs, spatial information, decision confidence, reward-predictive cues, and reward outcomes.^11–20^

Growing evidence indicates that the PFC contributes to a wide range of social behaviors, including social dominance and sexual behaviors.^21–24^ While most prior studies have focused on the medial prefrontal cortex (mPFC),^25^ our recent work in mice identified neural representations of pup retrieval, a hallmark maternal behaviors,^26^ in layer 5 OFC pyramidal neurons expressing the *Rbp4* gene^27^ (hereafter referred to as OFC^Rbp4^ neurons). These neurons constitute a major class of output cells projecting to subcortical targets.^28,29^ Maternal behaviors combine innate drives with experience-dependent flexibility: naïve virgin female mice can acquire these behaviors after repeated exposure to pups or observation of experienced mothers, underscoring their intrinsic learning capacity.^30,31^ This behavioral plasticity is shaped by neuromodulatory systems, including oxytocin and dopamine signaling.^27,31,32^ Silencing OFC output reduces the behaviorally locked activity of midbrain dopamine neurons during pup retrieval and delays the acquisition of maternal behaviors.^27^ However, the specific presynaptic partners of OFC^Rbp4^ neurons that support the formation and modulation of pup retrieval-related representations remain unknown.

Thalamocortical projections to the PFC have been implicated in cognitive flexibility, reversal learning in abstract tasks, and the maintenance of neural representations.^33–37^ They are also associated with sensory processing, attention, decision-making, and memory.^38^ Classical anatomical studies suggest that the OFC integrates inputs from multiple sensory cortices and higher-order thalamic nuclei, including the mediodorsal thalamus (MD) and the submedius thalamus (SMT).^39–41^ While the MD has been highlighted as a critical hub for cognitive flexibility,^33,34,37^ far less is known about the physiological functions of SMT, although its projections to OFC have been implicated in nociceptive modulation and goal-directed behavioral learning.^42–46^ Importantly, the contribution of these thalamic inputs to the neural representation of OFC during social behaviors remains poorly understood, partly due to limited genetic tools for selectively targeting specific neural subpopulations.

Recent advances in cell-type-specific genetic tools and high-resolution functional imaging now make it possible to dissect these circuits with unprecedented precision, particularly in the context of complex social behaviors, including maternal behaviors. Building on these methods, we systematically investigated the spatial organization and functional contribution of direct presynaptic input neurons to OFC^Rbp4^ neurons, with a particular focus on thalamocortical afferents.

## RESULTS

### Submedius thalamic nucleus is a prominent input source to the OFC

Our previous study demonstrated that OFC^Rbp4^ neurons shape neural representations during pup retrieval, thereby facilitating maternal behavioral learning.^27^ Mapping their presynaptic inputs is a critical first step toward elucidating the circuit mechanisms underlying the formation and modulation of these representations. To this end, we utilized Cre-dependent rabies virus-based retrograde monosynaptic tracing^47^ in *Rbp4-Cre* mice (Figure 1A). The majority of the starter cells, defined by colocalization of TVA-mCherry and rabies-derived GFP, were located in the OFC, predominantly within its ventral and lateral subdivisions (Figure 1B, C), which were previously identified as the primary loci of pup retrieval-related neural representations.^27^ To quantitatively analyze the distribution of presynaptic partners, we used AMaSiNe,^48^ a section-based whole-brain analysis platform that enables three-dimensional reconstruction and annotation of brain regions containing labeled neurons (Methods). Presynaptic neurons were primarily found in both ipsilateral and contralateral cortices and the thalamus, with smaller contributions from olfactory areas, the hippocampal formation, and the pallidum (Figure 1D–F; Figure S1).

**Figure 1.**
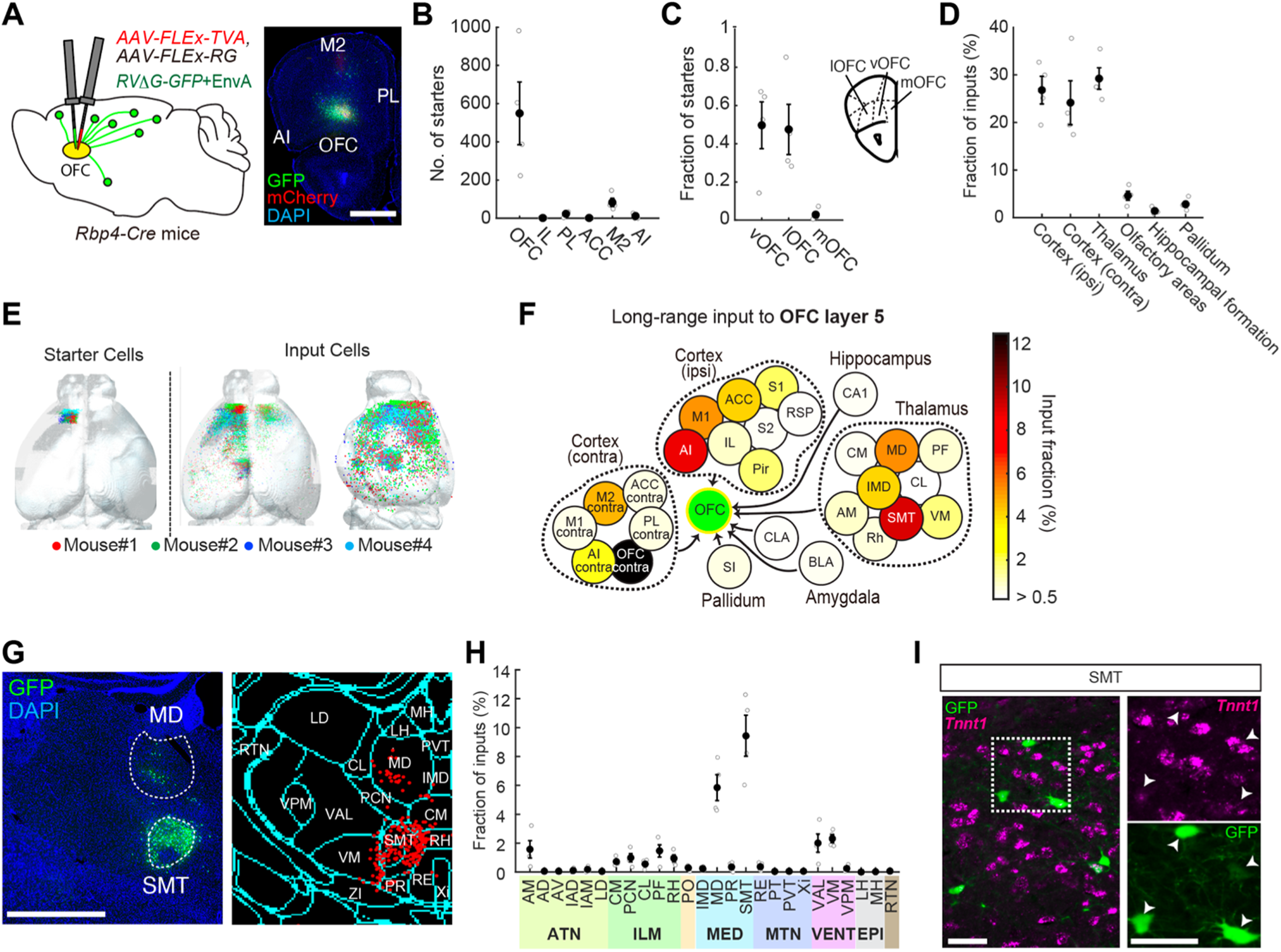
Presynaptic landscape of OFC^Rbp4^ neurons. (A) Experimental design (left) and a representative fluorescent micrograph of the injection site (right). Scale bar, 1 mm. (B) Number of starter cells in the frontal cortical areas; the majority are located in the OFC. (C) Fraction of starter cells within the OFC. Starter cells are located in the ventral and lateral OFC (vOFC and lOFC) but not in the medial OFC (mOFC). (D) Fraction of input cells across major brain regions. A large fraction is located bilaterally in the cortex and ipsilaterally in the thalamus (N = 4 mice). (E) Three-dimensional illustration of the anatomical distribution of the starter (left) and rabies-GFP-labeled input cells (middle and right). Data from four animals are superimposed on a common brain atlas. (F) Schematic map of selected long-range monosynaptic inputs to the OFC^Rbp4^ neurons. Colors indicate relative contribution of each region to the total input. Injection sites are shown by a green circle. (G) Left: Representative micrograph showing input neurons located in the MD and SMT. Scale bar, 1 mm. Right: Schematic of detected input cells and annotated thalamic regions using AMaSiNe. (H) Quantification of input cell proportions across thalamic regions. A large fraction originates from the SMT and MD. (I) Representative micrograph of rabies-GFP-labeled input neurons (green, detected using antibody staining) in the thalamus co-stained with a *Tnnt1* RNA probe using *in situ* hybridization. Scale bar, 50 μm. In total, 54 of 65 GFP+ cells were *Tnnt1*-positive. Error bars, SEM. See Figure S1 for more data. For abbreviations, see Table S1.

Because thalamocortical afferents are well suited for conveying pup-related sensorimotor signals to the OFC, we focused on thalamic inputs. Quantitative analyses revealed particularly prominent input from the submedius thalamus (SMT), in addition to the MD (Figure 1F–H; SMT: 9.4 ± 1.4%; MD: 5.8 ± 0.9% [mean ± standard error of the mean]). Notably, recent projection-based transcriptomic profiling identified marker genes specific to thalamic subdivisions.^49^ Among these, we found that the majority (83%) of rabies-GFP-expressing cells in the SMT were positive for *Tnnt1*, which encodes a subunit of troponin^49^ (Figure 1I). Thus, *Tnnt1* provides a useful marker for visualizing, monitoring, and manipulating SMT neurons projecting to the OFC.

### SMT and MD complimentary project to the frontal cortex

To achieve genetic access to *Tnnt1* neurons for anatomical and functional characterization of SMT, we generated *Tnnt1-Cre* knock-in mice using CRISPR-Cas9-mediated genome editing (Figure 2A, B). Crossing *Tnnt1-Cre* with *Ai162* reporter mice, which drive Cre-dependent GCaMP6s expression, yielded selective GCaMP6s labeling in the SMT and MD, along with several additional thalamic nuclei, consistent with reported mRNA expression patterns (Figure 2A).^49^ The vast majority of GCaMP6s-expressing neurons in both SMT and MD expressed *Tnnt1* and *vGluT2* mRNAs (Figure 2C–E), confirming that *Tnnt1-Cre* mice allow selective targeting of excitatory *Tnnt1* neurons in these regions (SMT^Tnnt1^ and MD^Tnnt1^ neurons, respectively) without affecting neighboring *Tnnt1*-negative cells.

**Figure 2.**
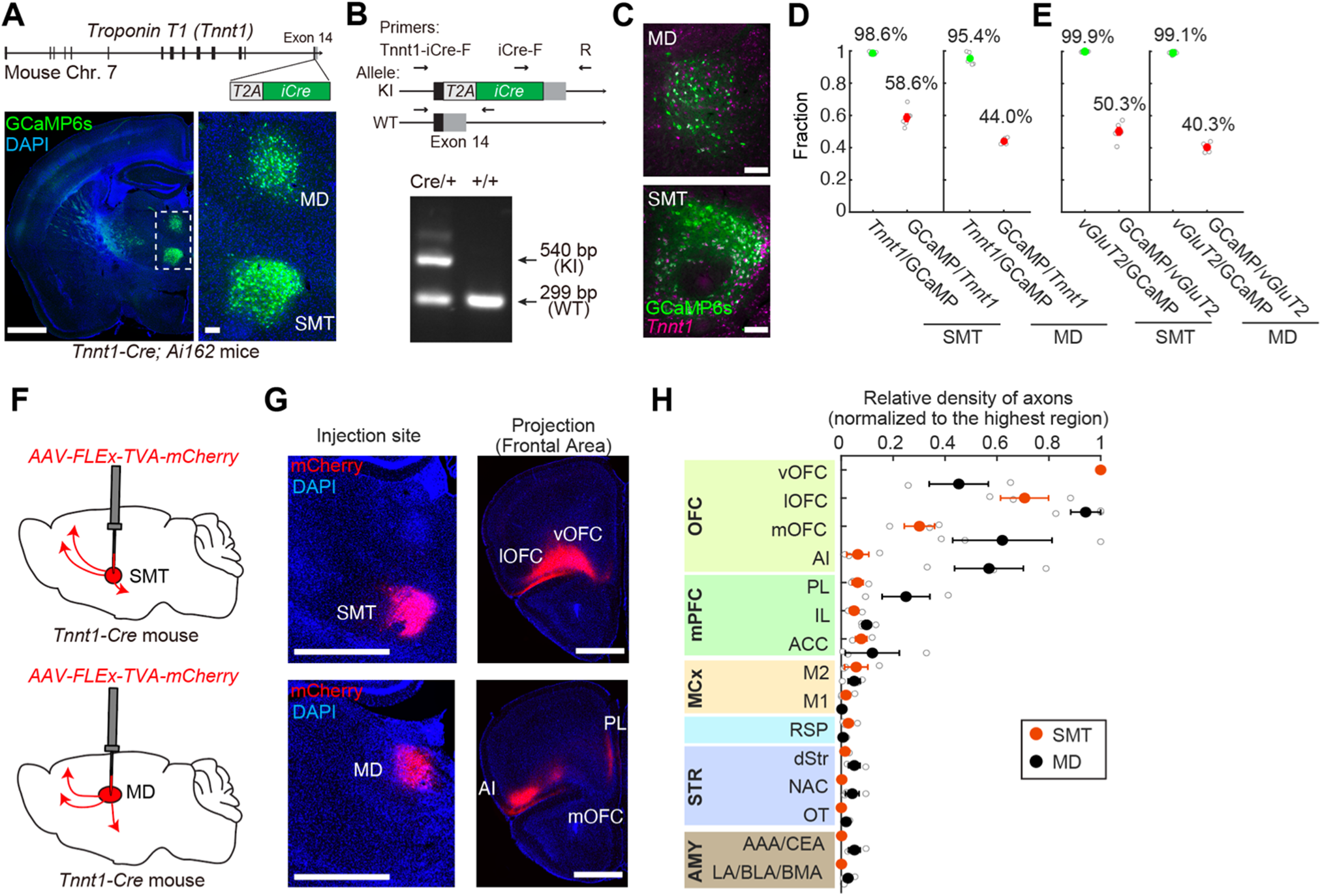
Complementary projection patterns of SMT and MD to the prefrontal cortex. (A) Top: Schematic of *Tnnt1-iCre* knock-in (KI) allele. Bottom: Representative micrograph of a *Tnnt1-Cre; Ai162* mouse (left) and a high-magnification view of reporter expression in the SMT and MD (right). Scale bar, 100 μm. (B) Top: Schematic of the primer design. Bottom: Representative electrophoresis image validating the iCre allele and negative control (wild-type [WT]). (C) Representative coronal sections showing *Tnnt1* mRNA expression (magenta) and GCaMP6s (anti-GFP, green) in the MD (top) and SMT (bottom) of a *Tnnt1-Cre; Ai162* mouse. (D, E) Quantification of specificity (*Tnnt1*+/GCaMP6s+ in panel D; *vGluT2*+/GCaMP6s+ in panel E) and efficiency (GCaMP6s+/*Tnnt1*+ in panel D; GCaMP6s+/*vGluT2*+ in panel E) (N = 6 hemispheres). GCaMP6s expression is restricted to *Tnnt1*+ neurons, whereas only a subset of *Tnnt1*+ neurons express GCaMP6s. The reduced efficiency may reflect stochastic inactivation of the *tetracycline responsive element* promoter in the *Ai162* line. A similar subset expression pattern has been reported in oxytocin (OT) neurons by *OT-Cre*; *Ai162* mice.^56^ (F) Schematic of the experimental design for axonal tracing. (G) Representative micrographs of injection sites (left) showing mCherry expression in the SMT (top) and MD (bottom). SMT axons primarily project to the ventral and lateral OFC (vOFC and lOFC) (right), whereas MD axons preferentially target the PL, medial OFC (mOFC), and AI. (H) Quantification of the axonal projection density from SMT^Tnnt1^ and MD^Tnnt1^ neurons across target regions (N = 3 mice). Error bars, SEM. See Figure S2 for more data. For abbreviations, see Table S1.

To map output pathways, we injected AAV *FLEx-TVA-mCherry*, which drives membrane-bound fluorescent labeling, into the SMT or MD of *Tnnt1-Cre* mice (Figure 2F). The number of mCherry-expressing neurons was quantified for each animal and used to normalize axonal projection strength (Figure S2A, B). Within the frontal cortex, SMT^Tnnt1^ neurons projected primarily to the ventral and lateral OFC (vOFC and lOFC), whereas MD^Tnnt1^ neurons projected more broadly to the medial and lateral OFC (mOFC and lOFC) as well as the agranular insular cortex (AI) (Figure 2G, H). In addition, MD^Tnnt1^, but not SMT^Tnnt1^, neurons sent axons to mPFC, including the prelimbic cortex (PL), as well as the amygdala and ventral striatum (Figure 2H). Thus, SMT^Tnnt1^ neurons selectively target OFC subregions previously implicated in pup retrieval-related neural representations, while MD^Tnnt1^ neurons provide broader and complementary projections to multiple prefrontal and limbic structures.

### Response properties of SMT during pup-retrieval

To examine the functional role of SMT in shaping pup retrieval representations in OFC^Rbp4^ neurons, we conducted microendoscopic Ca^2+^ imaging of SMT^Tnnt1^ neurons during pup retrieval in *Tnnt1-Cre; Ai162* female mice. A GRIN lens was implanted above the SMT, enabling neural recordings while virgin females engaged in pup retrieval after co-housing with lactating mothers (Figure 3A). To compare SMT^Tnnt1^ neuron activity across different stages of behavioral acquisition, we defined the first day on which a virgin retrieved more than ten pups across two 6-min imaging sessions as alloparental day 1 (AP1). On AP1, the number of retrievals was relatively low, but this parameter increased significantly on the subsequent day of co-housing (AP2) (Figure S3B). A representative AP2 dataset is shown in Figure 3B–D, with 60 regions of interest (ROIs; putatively single cells) identified in the field of view.

**Figure 3.**
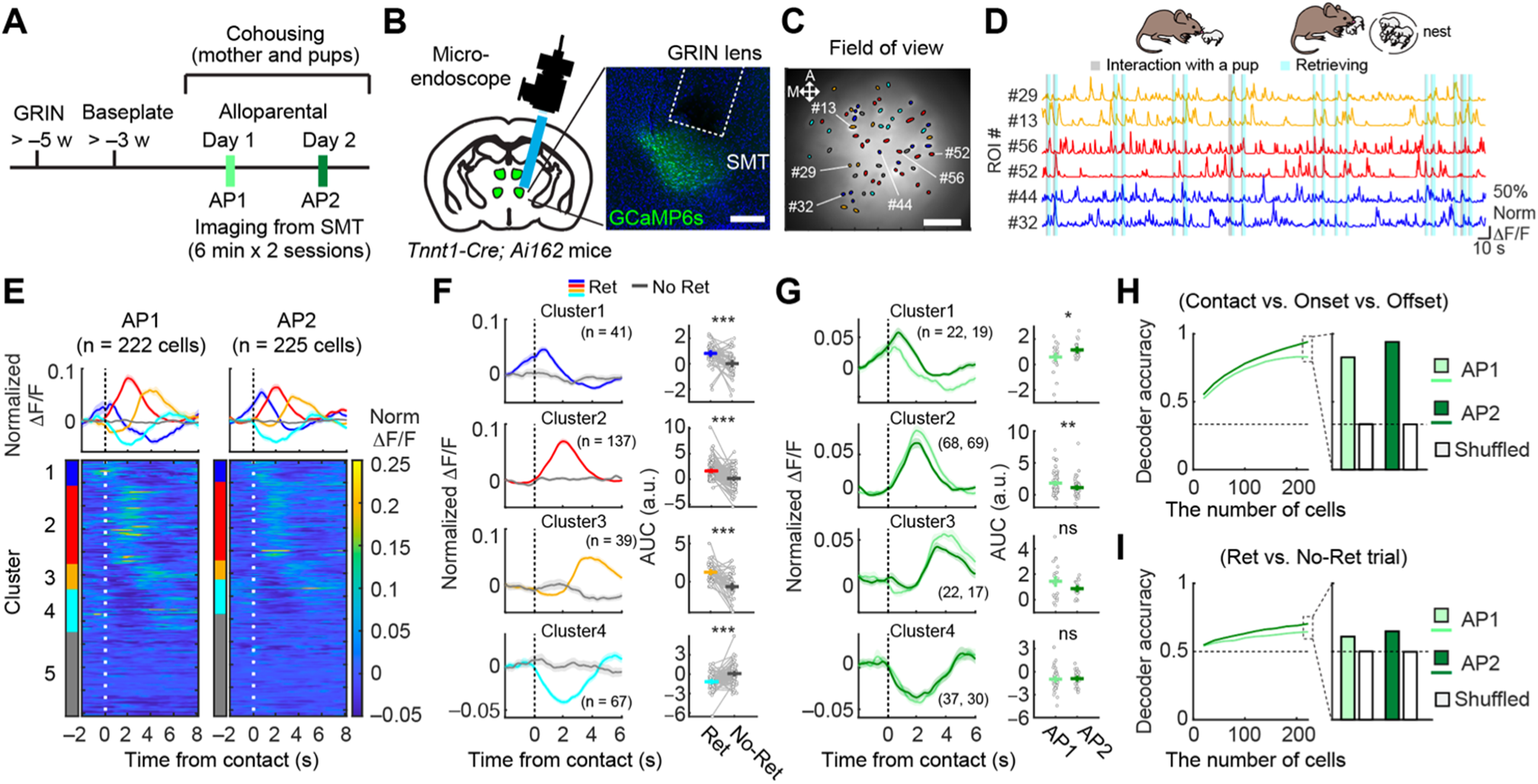
Neural representations of pup retrieval in SMT^Tnnt1^ neurons. (A) Schematic of the experimental design. (B) Left: Schematic of the head-attached microendoscope and GRIN lens implantation above the SMT. Right: Representative images showing the GRIN lens tract and GCaMP6s expression in *Tnnt1-Cre; Ai162* mice. Scale bar, 200 μm. (C) Spatial map of retrieval-responsive ROIs in a female mouse on AP2. ROIs are outlined in black, with colors indicating clusters (see Figure S3). A, anterior; M, medial. Scale bar, 100 μm. (D) Representative Ca^2+^ traces from ROIs belonging to Cluster 1 (blue, early), Cluster 2 (red, middle), and Cluster 3 (yellow, late). The indicated IDs correspond to the numbers in panel C. (E) Heat maps showing normalized trial-averaged responses during retrieval, sorted by cluster. Time 0 denotes pup contact followed by retrieval. (F) Left: Trial-averaged traces aligned to pup contact followed by retrieval (colored) or non-retrieval (gray). Right: AUC of PETHs. Numbers in parentheses indicate the number of ROIs per cluster (data pooled from AP1 and AP2). ***, p < 0.001 using the permutation test. (G) Left: Response activity during pup retrieval for each cluster. Right: AUC of PETHs. *, p < 0.05 and **, p < 0.01 using the permutation test. Numbers in parentheses indicate the number of ROIs in AP1 and AP2, respectively. (H, I) SVM decoding accuracy for classifying contact, onset, and offset of retrieval (H) and for classifying retrieval versus non-retrieval trials (I). Left: Accuracy as a function of cell number used for the training. Right: Accuracy using 220 cells. Dotted line indicates chance. Error bars, SEM. See Figure S3 for more data.

Pup retrieval unfolds in three sequential behavioral categories: pup contact, retrieval onset, and completion (placing the pup into the nest). To classify heterogeneous SMT^Tnnt1^ responses across AP1 and AP2, we conducted unbiased clustering of event-averaged Ca^2+^ signals. This analysis revealed four distinct response clusters, along with a non-responsive cluster (Figure S3C, D). These were qualitatively categorized as early (Cluster 1), middle (Cluster 2), late (Cluster 3), suppressed (Cluster 4), and non-responsive (Cluster 5). Heatmaps of cluster-sorted ROIs aligned to pup contact demonstrated that SMT^Tnnt1^ neurons collectively shaped the temporal structure of pup retrieval (Figure 3E). Across clusters, retrieval trials elicited significantly greater responses than non-retrieval trials (Figure 3F).

To assess experience-dependent changes, we calculated the area under the curve (AUC) of peri-event time histograms (PETHs) from each cluster. Cluster 1 neurons significantly increased activity from AP1 to AP2, whereas Cluster 2 neurons decreased activity (Figure 3G), indicating selective tuning of specific SMT^Tnnt1^ subsets during the acquisition of alloparental retrieval behavior. The relative proportions of cells in each cluster did not significantly differ between AP1 and AP2 (Figure S3E), suggesting that learning-related changes reflect tuning within existing ensembles rather than recruitment of new cells.

At the population level, we evaluated representational capacity using machine learning-based decoding analyses across AP1 and AP2. A linear support vector machine (SVM) trained to classify behavioral stages during pup retrieval (Methods) showed increasing accuracy with larger training sets of ROIs, with higher decoding accuracy at AP2 than AP1. Shuffling the dataset reduced classification accuracy to near-chance levels (Figure 3H). These results indicate that SMT^Tnnt1^ populations acquire greater representational precision with behavioral experience. Beyond encoding ongoing behavioral states, SMT activity during pup contact was sufficient to predict retrieval outcome, with decoding accuracy modestly higher at AP2 (Figure 3I).

In summary, behaviorally responsive SMT^Tnnt^^1^ neurons undergo experience-dependent refinement and encode sufficient information to classify behavioral stages and predict behavioral outcomes during pup retrieval.

### Response properties of MD during pup-retrieval

To compare physiological properties between SMT and MD, we conducted parallel equivalent microendoscopic Ca^2+^ imaging of MD^Tnnt1^ neurons during pup retrieval in *Tnnt1-Cre; Ai162* female mice (Figure 4A). A representative AP2 dataset is shown in Figure 4B–D, with 47 ROIs identified within the field of view. Unbiased clustering analysis revealed four distinct response clusters and one non-responsive cluster in MD^Tnnt1^ neurons, qualitatively similar to SMT (Figure 4E; Figure S3E). Heatmap visualization demonstrated that MD^Tnnt1^ neurons, as a population, also represented the temporal structure of pup retrieval, with activity patterns resembling those observed in SMT (Figure 4E). Across all responsive clusters, MD^Tnnt1^ neurons displayed significant transient activity during pup retrieval compared with non-retrieval, confirming MD’s active engagement in this behavior (Figure 4F).

**Figure 4.**
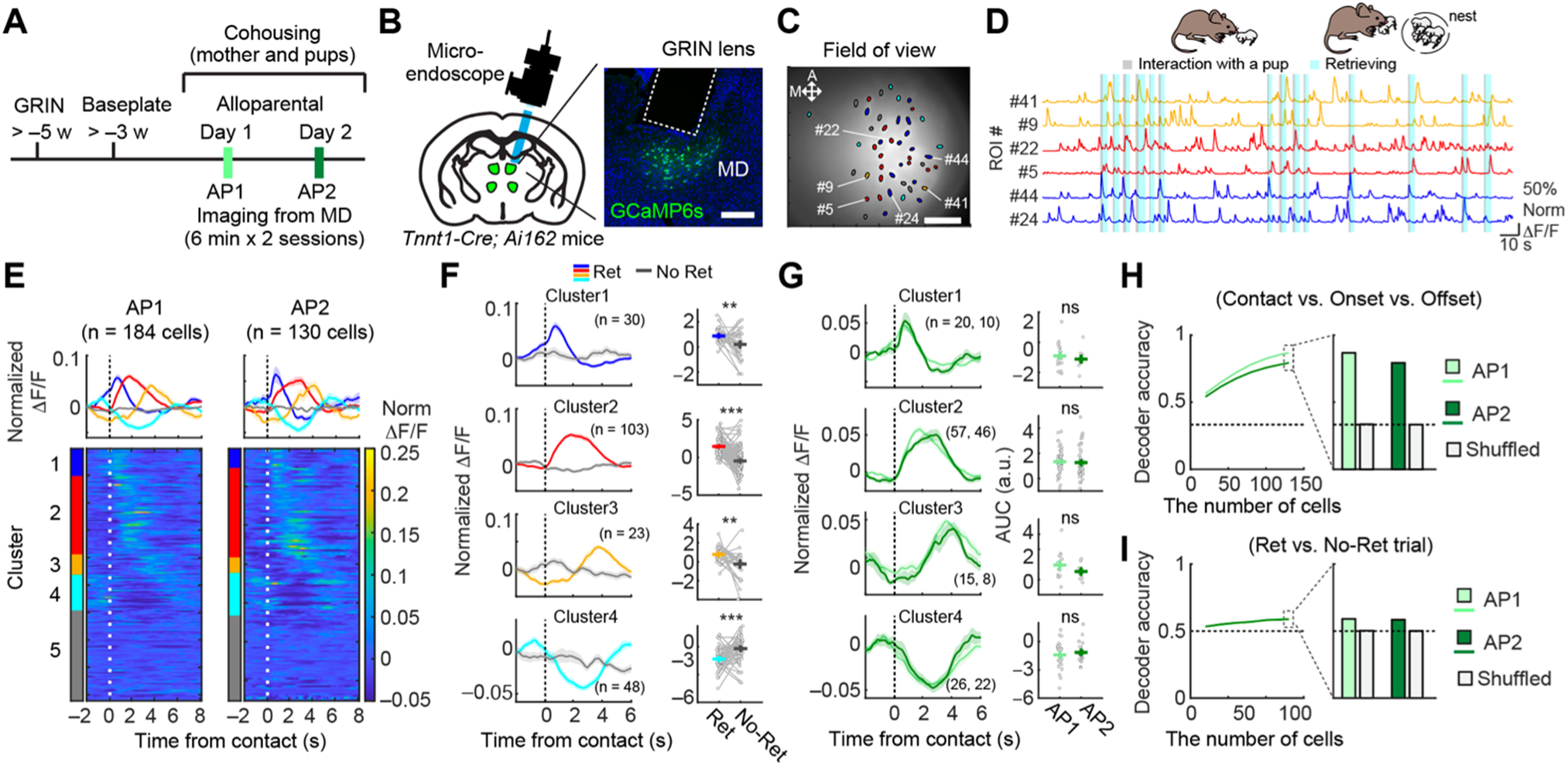
Neural representations of pup retrieval in MD^Tnnt1^ neurons. (A) Schematic of the experimental design. (B) Left: Schematic of the head-attached microendoscope and GRIN lens implantation above the MD. Right: Representative images showing the GRIN lens tract and GCaMP6s expression in *Tnnt1-Cre; Ai162* mice. Scale bar, 200 μm. (C) Spatial map of retrieval-responsive ROIs in a female mouse on AP2. ROIs are outlined in black, with colors indicating clusters. A, anterior; M, medial. Scale bar, 100 μm. (D) Representative Ca^2+^ traces from ROIs belonging to Cluster 1 (blue, early), Cluster 2 (red, middle), and Cluster 3 (yellow, late). The indicated IDs correspond to the numbers in panel C. (E) Heat maps showing normalized trial-averaged responses during retrieval, sorted by cluster. Time 0 denotes pup contact followed by retrieval. (F) Left: Trial-averaged traces aligned to pup contact followed by retrieval (colored) or non-retrieval (gray). Right: AUC of PETHs. Numbers in parentheses indicate the number of ROIs per cluster (data pooled from AP1 and AP2). ***, p < 0.001 using the permutation test. (G) Left: Response activity during pup retrieval for each cluster. Right: AUC of PETHs. *, p < 0.05 and **, p < 0.01 using the permutation test. Numbers in parentheses indicate the number of ROIs in AP1 and AP2, respectively. (H, I) SVM decoding accuracy for classifying contact, onset, and offset of retrieval (H) and for classifying retrieval versus non-retrieval trials (I). Left: Accuracy as a function of cell number used for the training. Right: Accuracy using 130 cells. Dotted line indicates chance. Error bars, SEM. See Figure S3 for more data.

A key difference emerged when examining learning-related plasticity. In contrast to SMT, MD^Tnnt1^ neurons showed stable activity patterns across all clusters between AP1 and AP2 (Figure 4G). AUC analysis of PETHs revealed no significant changes, indicating that MD response properties remain invariant across behavioral stages. This stability suggests distinct roles for SMT and MD: whereas SMT undergoes experience-dependent tuning, MD provides stable baseline representations.

Population-level decoding analyses further reinforced this dissociation. MD^Tnnt1^ neurons reliably classified behavioral categories during pup retrieval, and decoding accuracy increased with larger ROI sets. However, unlike SMT, decoding performance did not improve from AP1 to AP2 (Figure 4H). Similarly, MD activity during the pup-contact window only modestly predicted retrieval outcomes, with decoding accuracy unchanged across learning stages (Figure 4I). Together, these results demonstrate that both SMT^Tnnt1^ and MD^Tnnt1^ neurons encode behaviorally relevant information during pup retrieval. However, only SMT exhibits experience-dependent refinement, enhancing representational precision and predictive capacity as maternal behavior acquisition progresses. In contrast, MD provides stable, learning-invariant representations that may serve as a consistent reference signal for maternal behavior control.

### SMT contributes to shaping pup-retrieval representations in OFC^Rbp^^4^ neurons

To directly assess the functional contributions of SMT and MD to pup retrieval representations in OFC^Rbp4^ neurons, we combined projection-specific chemogenetic inhibition with microendoscopic Ca^2+^ imaging of OFC^Rbp4^ neurons in *Rbp4-Cre; Ai162* female mice (Figure 5A). To selectively target OFC-projecting neurons in SMT or MD (SMT^→OFC^ or MD^→OFC^), we injected AAV-retro *EF1a-FlpO* into the OFC and a FlpO-dependent hM4Di vector (AAV *hSyn-fDIO-hM4Di-mCherry*) bilaterally into SMT or MD, 2–3 weeks before GRIN lens implantation above the OFC (Figure 5A, B; Figure S4A, B). This approach enabled causal testing of whether the anatomical connections identified translate into functional influences on OFC activity during pup retrieval.

**Figure 5.**
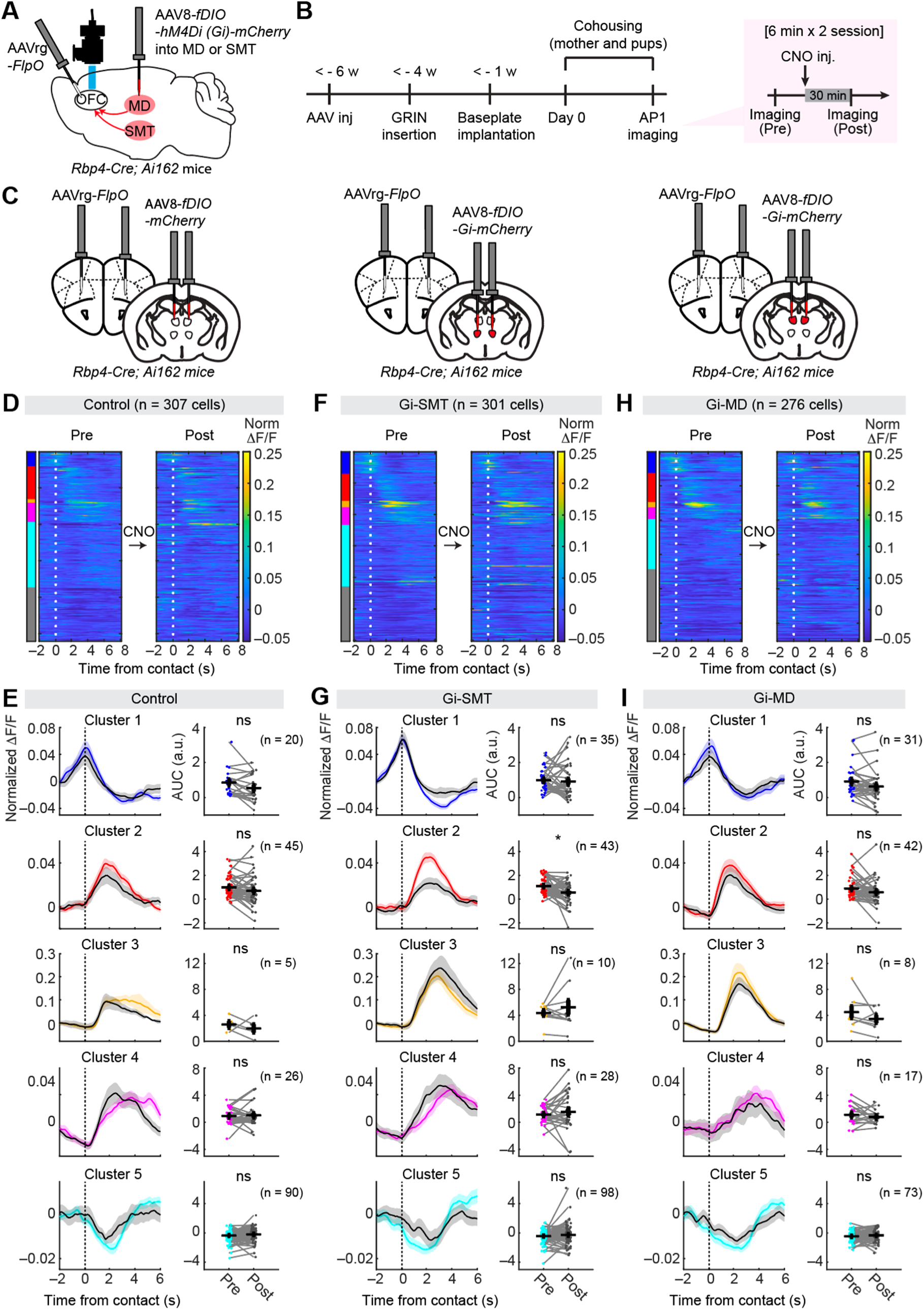
SMT functionally modulates pup-retrieval representations in the OFC^Rbp4^ neurons. (A) Schematic of the experimental design. (B) Timeline of the experimental paradigm. Two 6-min imaging sessions were performed before and after CNO administration. (C) Schematic of the virus injection strategy. (D, F, H) Heat maps of normalized, trial-averaged single-cell responses during pup retrieval before (left) and after (right) CNO. Time 0 denotes pup contact followed by retrieval. (E, G, I) Quantification of averaged responses before and after CNO. *, p < 0.05 using the permutation test. ns, not significant. Numbers in parentheses indicate the number of ROIs per cluster. See Figure S4 for more data.

We performed Ca^2+^ imaging during pup retrieval both before and after CNO administration, enabling within-subject comparisons at the single-cell level. Negative controls expressed mCherry alone (Figure 5C). Unbiased clustering analysis revealed five response clusters and one non-responsive cluster in OFC^Rbp4^ neurons, consistent with our previous findings^27^ (Figure S4D). Control experiments confirmed that CNO injection alone did not alter behavioral performance (Figure S4C) or neural representation patterns across clusters (Figure 5D, E). Moreover, silencing either SMT^→OFC^ or MD^→OFC^ neurons did not impair pup retrieval behavior (Figure S4C), allowing us to isolate circuit-level effects on OFC representations from behavioral confounds.

Silencing SMT^→OFC^ neurons selectively attenuated Cluster 2 responses in the OFC, corresponding to neurons active during the middle phase of pup retrieval sequences (Figure 5F, G). Other responsive clusters were unaffected, indicating that SMT provides temporally specific modulation of distinct OFC subpopulations. In contrast, silencing MD^→OFC^ neurons produced no detectable changes across any responsive clusters (Figure 5H, I). This lack of effect, despite substantial anatomical connectivity, suggests that MD projections to OFC do not significantly contribute to maternal behavior representations under these conditions. In summary, SMT, but not MD, exerts selective and functionally specific influence on pup retrieval representations in OFC^Rbp4^ neurons.

## DISCUSSION

The OFC is important for diverse cognitive functions and social behaviors, yet its precise presynaptic partners and their functional contributions remain incompletely defined. Here, we provide a systematic functional characterization of higher-order thalamic inputs to the OFC in the context of maternal behaviors in female mice, identifying the SMT as a previously unrecognized node of functional relevance. Below, we summarize the key insights from this study and acknowledge its limitations.

### SMT as a presynaptic node of OFC in maternal behavior circuitry

Rabies virus-based tracing identified SMT as a major source of thalamocortical afferents to layer 5 excitatory neurons in the OFC, consistent with findings from previous classical tracer studies.^41,50–53^ Using a newly developed and validated Cre-driver mouse line, we demonstrate that SMT^Tnnt1^ neurons robustly encode pup retrieval behavior, exhibiting both anticipatory activity preceding pup contact and responses to ongoing motor sequences. With behavioral experience, SMT neurons exhibited enhanced anticipatory responses and diminished retrieval onset-related responses, suggesting learning-related plasticity. These changes may reflect the integration of predictive signals about pup-directed actions, potentially facilitating more efficient and fluid behavioral sequences. Notably, such plasticity was absent in MD neurons despite superficially similar activity patterns, underscoring functional specialization among higher-order thalamocortical circuits.^45^ Moreover, only SMT^Tnnt1^, and not MD^Tnnt1^, neurons exerted functional influence over OFC^Rbp4^ neuron activity. Consistent with this dissociation, MD displayed relatively inferior decoding performance of pup retrieval compared with SMT (compare Figure 3H–I with Figure 4H–I). Collectively, these findings suggest that thalamic nuclei subserve distinct computational roles in shaping complex, learned behaviors.

A key advance of this work is the generation of the *Tnnt1-Cre* mouse line, which enables unprecedented specificity for targeting higher-order thalamic neurons and overcomes technical challenges inherent to probing deep thalamic circuits. Using this tool, we demonstrated that SMT^Tnnt1^ and MD^Tnnt1^ neurons, although sharing molecular identity, display distinct projection topographies (Figure 2G, H). SMT projections were concentrated in lateral OFC regions, whereas MD projections extended more broadly, including medial prefrontal regions. This anatomical divergence likely underpins their differential contributions: SMT may exert focused influence on OFC circuits involved in action planning and behavioral flexibility,^45^ whereas MD may participate more broadly in prefrontal-dependent functions such as working memory and cognitive control.^33,34,37^ Future studies utilizing the *Tnnt1-Cre* line will be essential to dissect SMT and MD contributions across a wider range of cognitive and social tasks, clarifying how higher-order thalamic nuclei diversify cortical computations in adaptive behaviors.

### Potential circuit mechanisms and computational roles

How do SMT^Tnnt1^ neurons influence pup retrieval-related representations in OFC^Rbp4^ neurons? The temporal dynamics of their activity provide important clues. SMT^Tnnt1^ neurons exhibited relatively weaker and slower anticipatory responses (Cluster 1, Figure 3) compared to OFC^Rbp4^ neurons (Cluster 1, Figure 5), and silencing SMT^→OFC^ projectors did not disrupt these OFC anticipatory responses. These observations suggest that SMT inputs do not uniformly shape all OFC responses but instead exert more selective influence.

A possibility is that SMT neurons connect preferentially with a subset of OFC^Rbp4^ neurons, particularly those active at retrieval onset (Cluster 2, Figure 5). This selective connectivity would imply that SMT contributes to specific computational aspects of pup retrieval. Consistent with this view, both OFC^Rbp4^ neurons^27^ and SMT^Tnnt1^ neurons (Cluster 2, Figure 3) exhibit a parallel reduction in retrieval-onset responses between AP1 and AP2, suggesting functional coupling between these populations. A second possibility is that SMT operates downstream of OFC. Although our circuit-mapping emphasized the presynaptic nature of SMT inputs (Figure 1), SMT and OFC are reciprocally connected.^44,53,54^ In this reciprocal framework, SMT may serve to amplify and prolong behaviorally relevant representations initiated in OFC, sustaining them across the extended temporal structure of pup retrieval. A third possibility is that SMT and OFC act as components of a distributed, parallel-processing network, with neither structure serving as a simple upstream driver. Instead, each may encode partially distinct aspects of pup-directed behavior that converge to generate coherent maternal behavioral sequences. This interpretation is supported by the differential learning-related plasticity observed between SMT and OFC. While both regions encode pup retrieval, only SMT showed enhanced anticipatory responses (Cluster 1) in AP2 compared with AP1, suggesting specialized tuning to optimize predictive signals for upcoming maternal behaviors.

Distinguishing among these models will require simultaneous high-resolution recordings from SMT and OFC, combined with temporally precise optogenetic perturbations.^46^ Such approaches will be essential for determining the causal flow of information within this circuit and to understand how SMT contributions integrate with other presynaptic inputs to shape OFC representations of pup retrieval.

### Limitations of this study

Several limitations should be acknowledged. First, although chemogenetic silencing of SMT^→OFC^ neurons reduced OFC activity, the effect sizes were modest and did not produce overt behavioral deficits. This may reflect incomplete silencing achieved with projection-based targeting, functional compensation by parallel circuits, or the inherently distributed control of maternal behavior. Future studies using cell type–specific, temporally precise optogenetic perturbations will be necessary to establish stronger causal relationships. Second, we did not detect neural representations in the PFC that were clearly influenced by MD^Tnnt1^ neurons. Both SMT and MD neurons may contribute to pup retrieval through distinct PFC subnetworks, with SMT exerting more direct influence on the lOFC and vOFC regions targeted in our Ca^2+^ imaging experiments. Expanding analyses to additional PFC subregions will be critical for clarifying the role of MD in pup retrieval. Third, our manipulations targeted the effects of SMT and MD inputs on layer 5 OFC neurons. While this layer indeed receives substantial inputs from both regions, other cortical layers are also likely to integrate thalamic afferents^55^, which may play distinct roles in shaping OFC computations. Thus, future studies should systematically examine how laminar specificity in the OFC contributes to the functional impact of thalamocortical inputs. Fourth, our study focused exclusively on pup retrieval, which represents only one component of the broader maternal repertoire.^26^ Whether SMT contributions extend to other behaviors, such as nursing, grooming, or nest building, remains to be determined. Despite these limitations, the *Tnnt1-Cre* line, combined with microendoscopic imaging of SMT and MD neurons, provides a valuable platform for further dissecting how higher-order thalamic neurons regulate maternal and social behaviors in a broader behavioral spectrum.

## Resource Availability

### Lead contact

Further information and requests for resources and reagents should be directed to and will be fulfilled by the Lead Contact, Kazunari Miyamichi.

### Materials availability

All data supporting the findings of this study are available in the main paper and supplementary materials. All materials, including the *Tnnt1-Cre* mice, are available upon request from the corresponding authors.

### Data and code availability

- Original microendoscope data will be deposited in the SSBD repository at the time of publication.
- All custom MATLAB analysis scripts will be deposited in the SSBD repository at the time of publication.

## Acknowledgments

We thank members of the Miyamichi laboratory for the critical reading of the manuscript; Satsuki Irie for technical support; Edward M. Callaway for sharing the B7GG, BHK-EnvA, and HEK293-TVA800 cell lines; the University of North Carolina Vector Core and the Canadian Neurophotonics Platform Viral Vector Core Facility for AAV production; Masanori Murayama for providing the *Rbp4-Cre* mice; and the RIKEN BDR animal facility for animal care and *in vitro* fertilization; Jun Song and Se-Bum Paik for their technical consultation on the use of AMaSiNe. This work was supported by the JST PRESTO program (JPMJPR21S7), JSPS KAKENHI (20K15941 and 25K18590), RIKEN incentive and diversity promotion grants to G.T., and by a Takeda Science Foundation Research Grant, JSPS KAKENHI (21H02587 and 25K02368) to K.M.

## Author Contributions

G.T. and K.M. conceived the experiments. G.T. performed the experiments and analyzed the data, with technical support from M.H. and H.K. M.K. and T.A. designed and generated the *Tnnt1-Cre* mice, and M.H. confirmed their genomic structure. G.T. and K.M. drafted the manuscript with input from all authors.

## Supplementary Figures

**Figure S1.**
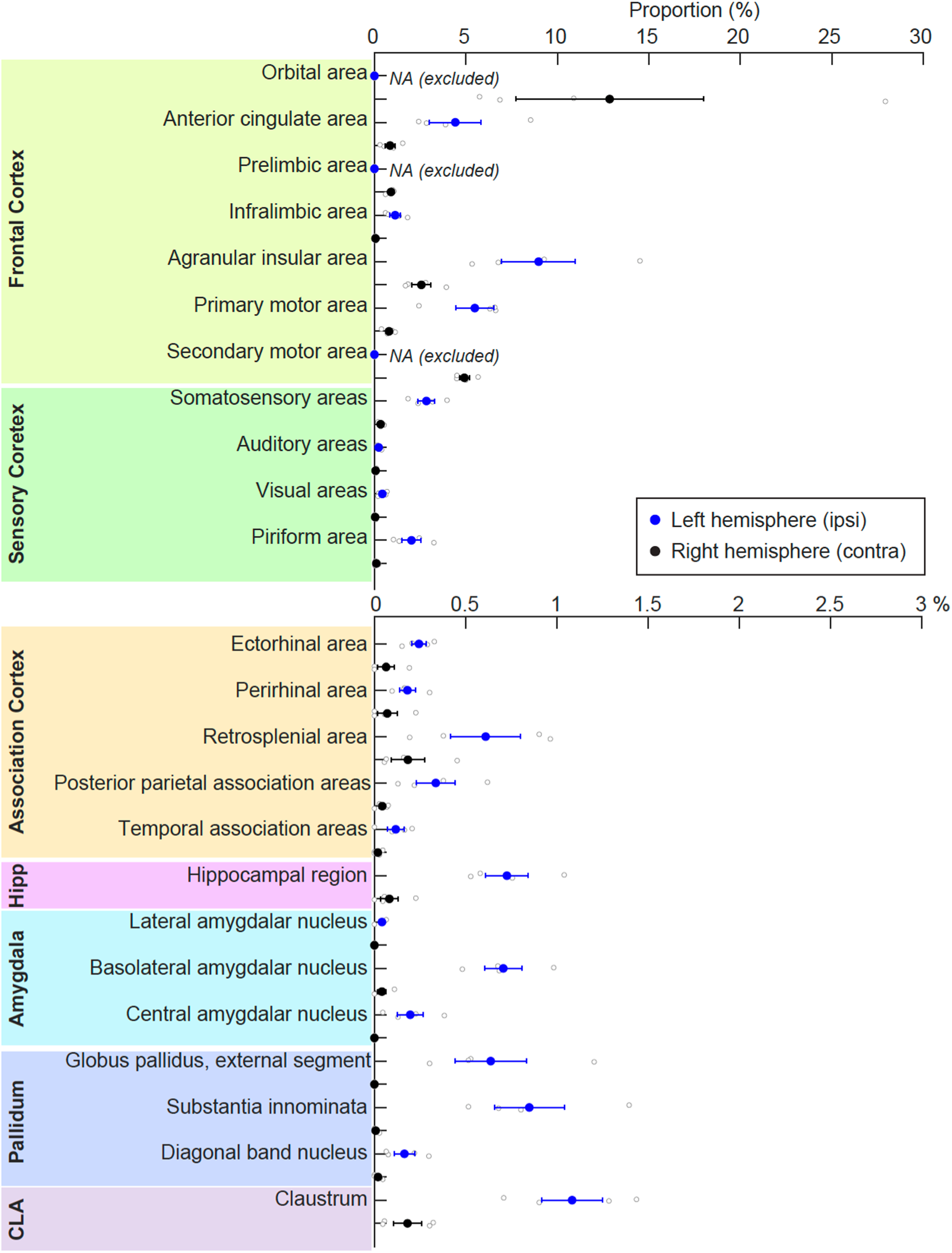
Whole-brain presynaptic distribution for OFC^Rbp4^ neurons, related to Figure 1. Quantification of rabies-GFP-labeled input cell proportions across cortical and subcortical areas (N = 4 mice). Error bars, SEM.

**Figure S2.**
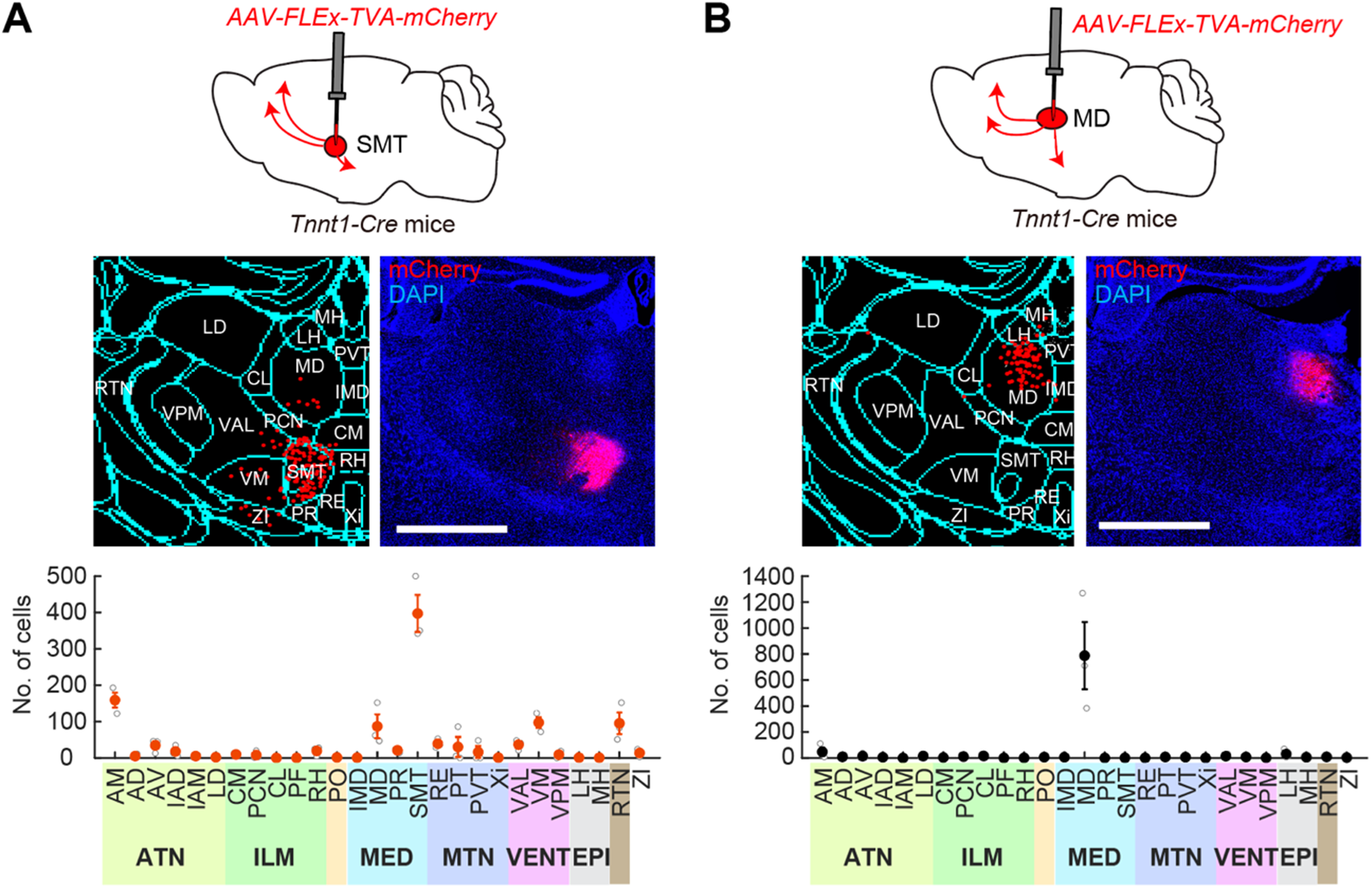
Regional distribution of labeled cells in *Tnnt1-Cre* axon tracing experiments, related to Figure 2. (A, B) Top: Schematic of the virus injection strategy. AAV5 *CAG-FLEx-TVA-mCherry* was injected into the left SMT (A) or MD (B). Middle left: Schematic of detected soma and annotated thalamic regions using AMaSiNe. Middle right: Representative coronal section of the injection site. Scale bar, 1 mm. Bottom: Quantification of the fraction of mCherry+ cells across thalamic areas (N = 3 mice).

**Figure S3.**
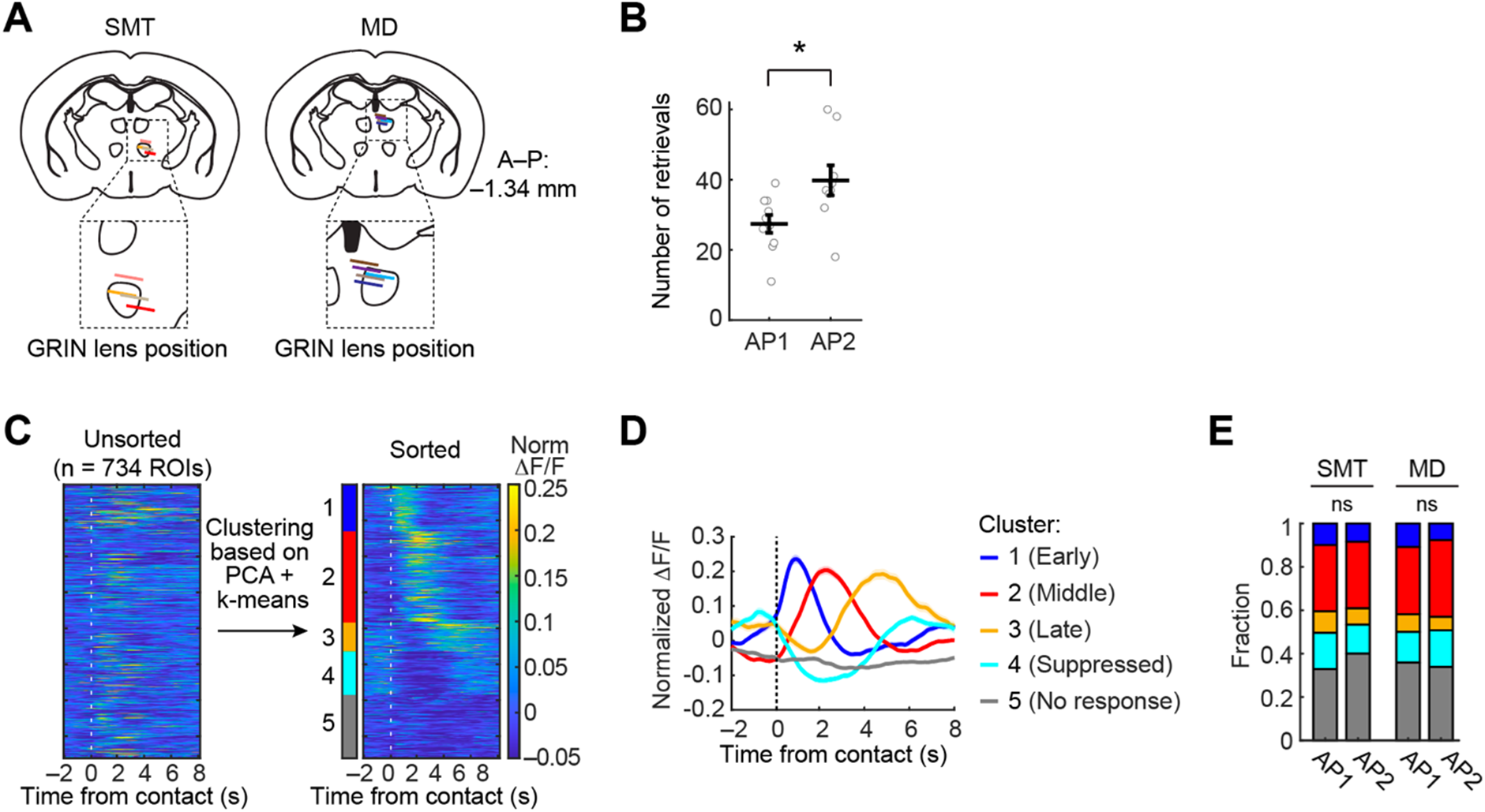
Clustering of SMT and MD ROIs based on Ca^2+^ responses during pup retrieval, related to Figures 3 and 4. (A) Schematics of coronal sections showing GRIN lens placement in the animals used for Figure 3 (Top, SMT) and Figure 4 (Bottom, MD). Lines denote the GRIN lens tip for each mouse (N = 4 for SMT and N = 5 for MD; one SMT animal was excluded because of death before sectioning). (B) Number of pup retrievals during imaging sessions. N = 10 and 9 mice for AP1 and AP2, respectively. *, p < 0.05 using the permutation test. (C) Normalized, averaged responses of ROIs during retrieval trials before (left) and after (right) clustering. (D) Trial-averaged normalized (norm) ΔF/F traces of ROIs by cluster. ΔF/F values were normalized to each ROI’s maximum. (E) Fraction of cells in each cluster. ns, not significant using the chi-square test.

**Figure S4.**
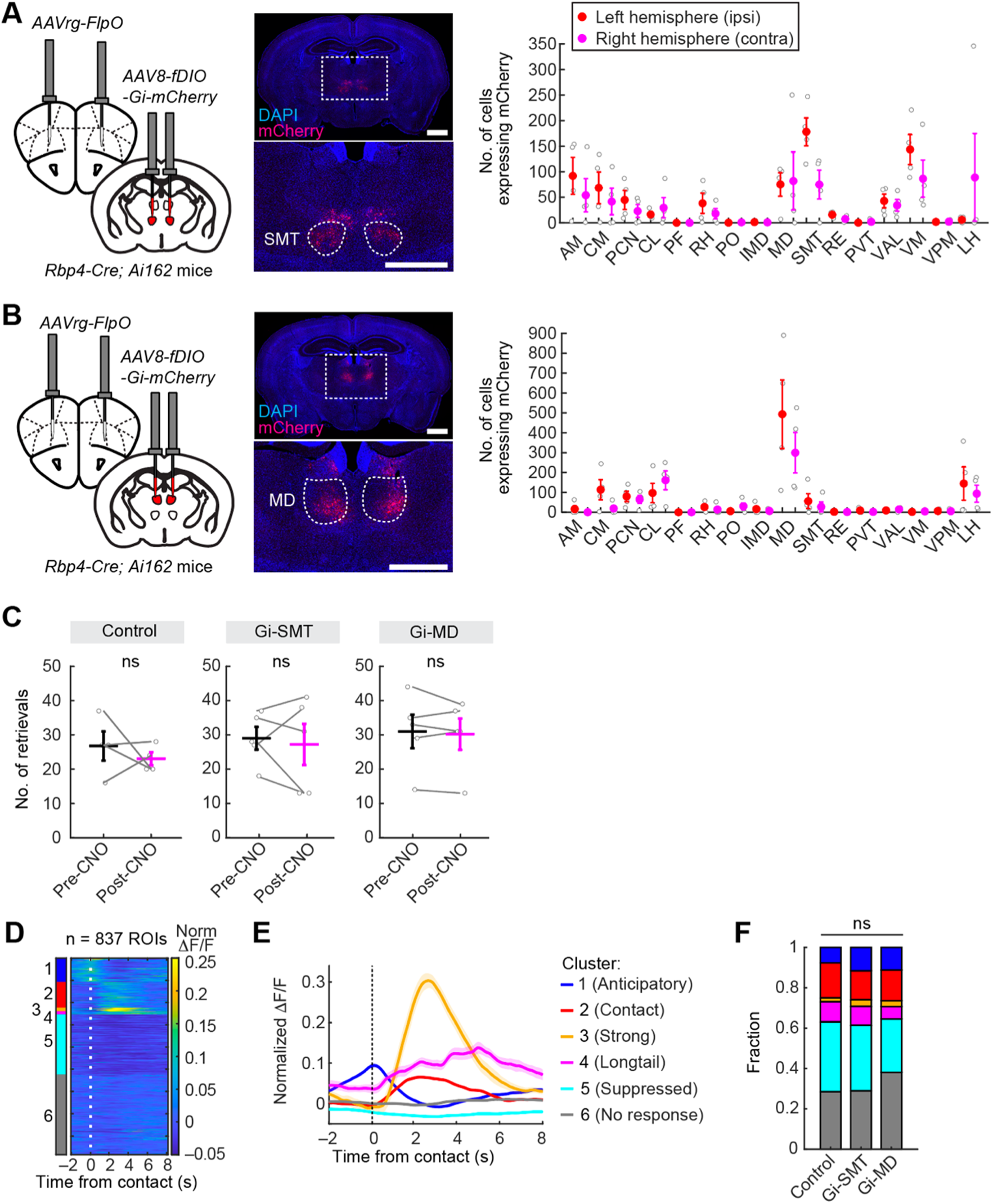
Chemogenetic silencing of SMT^→OFC^ or MD^→OFC^ neurons does not alter pup-retrieval performance, related to Figure 5. (A, B) Left: Schematic of the virus injection strategy. AAV8 *hSyn-fDIO-Gi-mCherry* was injected bilaterally into the SMT (A) or MD (B), and AAVrg *EF1a-FlpO* was injected bilaterally into the OFC. Middle: Representative coronal sections showing injection sites in the SMT (A) or MD (B). Scale bar, 1 mm. Right: Quantification of mCherry-expressing cells in the thalamus (N = 4 mice for panel A; N = 5 mice for panel B). (C) Number of pup retrievals during the imaging sessions. ns, not significant using the permutation test. Error bars, SEM. (D) Normalized, averaged responses of ROIs during retrieval trials after clustering. (E) Trial-averaged normalized (norm) ΔF/F traces of ROIs by cluster. ΔF/F values were normalized to each ROI’s maximum. (F) Fraction of cells within each cluster. ns, not significant using the chi-square test with Bonferroni correction.

**Table S1.** List of abbreviations, related to Figure 1–2.

| Acronym | Name |
| --- | --- |
| AAA | anterior amygdalar area |
| ACC | anterior cingulate cortex |
| AD | anterodorsal nucleus |
| AI | agranular insular cortex |
| AM | anteromedial nucleus |
| ATN | anterior group of the dorsal thalamus |
| AV | anteroventral nucleus |
| BLA | basolateral amygdalar nucleus |
| BMA | basomedial amygdalar nucleus |
| CA1 | hippocampus CA1 |
| CEA | central amygdalar nucleus |
| CL | central lateral nucleus |
| CLA | Clastrum |
| CM | central medial nucleus |
| dStr | dorsal striatum |
| EPI | Epithalamus |
| IAD | interanterodorsal nucleus |
| IAM | interanteromedial nucleus |
| IL | infralimbic cortex |
| ILM | intralaminar nuclei of the dorsal thalamus |
| IMD | intermediodorsal nucleus |
| LA | lateral amygdalar nucleus |
| LD | lateral dorsal nucleus |
| LH | lateral habenula |
| M1 | primary motor cortex |
| M2 | secondary motor cortex |
| MD | mediodorsal thalamus |
| MED | medial group of the dorsal thalamus |
| MH | medial habenula |
| MTN | midline group of the dorsal thalamus |
| NAC | nucleus accumbens |
| OT | olfactory tubercle |
| PCN | paracentral nucleus |
| PF | parafascicular nucleus |
| Pir | piriform cortex |
| PL | prelimbic cortex |
| PO | posterior complex of the thalamus |
| PR | perieuneus nucleus |
| PT | parataenial nucleus |
| PVT | paraventricular nucleus |
| RE | nucleus of reuniens |
| RH | rhomboid nucleus |
| RSP | retrosplenial cortex |
| RTN | reticular nucleus |
| S1 | primary somatosensory cortex |
| S2 | secondary somatosensory cortex |
| SI | substantia innominate |
| SMT | submedius thalamus |
| VAL | ventral anterior-lateral complex of the thalamus |
| VENT | ventral group of the dorsal thalamus |
| VM | ventral medial nucleus |
| VPM | ventral posteromedial nucleus |
| Xi | xiphoid thalamic nucleus |

## KEY RESOURCES TABLE

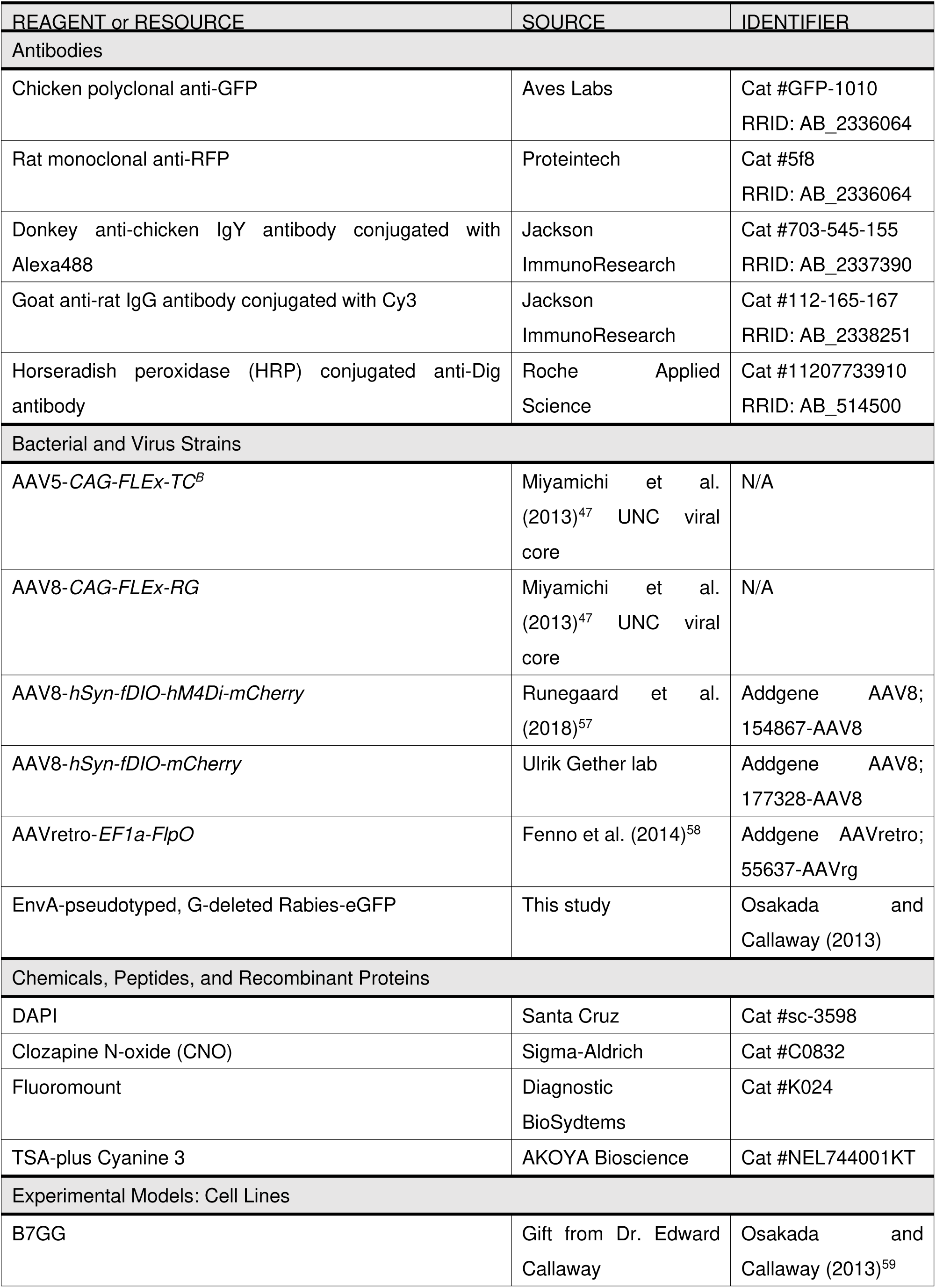
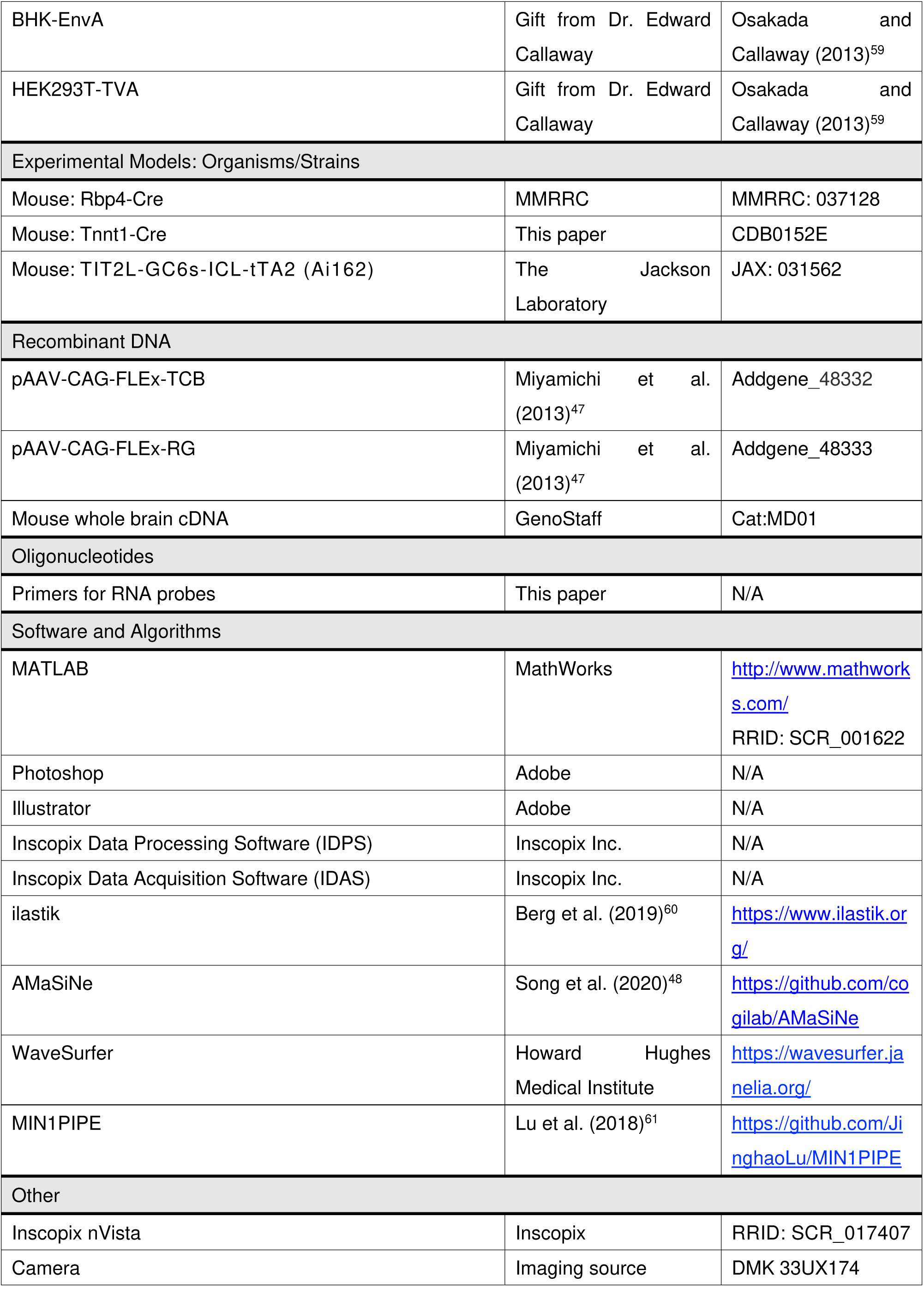

| REAGENT or RESOURCE | SOURCE | IDENTIFIER |
| --- | --- | --- |
| **Antibodies** |  |  |
| Chicken polyclonal anti-GFP | Aves Labs | Cat #GFP-1010<br>RRID: AB_2336064 |
| Rat monoclonal anti-RFP | Proteintech | Cat #5f8<br>RRID: AB_2336064 |
| Donkey anti-chicken IgY antibody conjugated with Alexa488 | Jackson ImmunoResearch | Cat #703-545-155<br>RRID: AB_2337390 |
| Goat anti-rat IgG antibody conjugated with Cy3 | Jackson ImmunoResearch | Cat #112-165-167<br>RRID: AB_2338251 |
| Horseradish peroxidase (HRP) conjugated anti-Dig antibody | Roche Applied Science | Cat #11207733910<br>RRID: AB_514500 |
| **Bacterial and Virus Strains** |  |  |
| AAV5-*CAG-FLEX-TC<sup>B</sup>* | Miyamichi et al. (2013)<sup>47</sup> UNC viral core | N/A |
| AAV8-*CAG-FLEX-RG* | Miyamichi et al. (2013)<sup>47</sup> UNC viral core | N/A |
| AAV8-*hSyn-fDIO-hM4Di-mCherry* | Runegaard et al. (2018)<sup>57</sup> | Addgene AAV8; 154867-AAV8 |
| AAV8-*hSyn-fDIO-mCherry* | Ulrik Gether lab | Addgene AAV8; 177328-AAV8 |
| AAVretro-*EF1a-FlpO* | Fenno et al. (2014)<sup>58</sup> | Addgene AAVretro; 55637-AAVrg |
| EnvA-pseudotyped, G-deleted Rabies-eGFP | This study | Osakada and Callaway (2013) |
| **Chemicals, Peptides, and Recombinant Proteins** |  |  |
| DAPI | Santa Cruz | Cat #sc-3598 |
| Clozapine N-oxide (CNO) | Sigma-Aldrich | Cat #C0832 |
| Fluoromount | Diagnostic BioSystems | Cat #K024 |
| TSA-plus Cyanine 3 | AKOYA Bioscience | Cat #NEL744001KT |
| **Experimental Models: Cell Lines** |  |  |
| B7GG | Gift from Dr. Edward Callaway | Osakada and Callaway (2013)<sup>59</sup> |
| BHK-EnvA | Gift from Dr. Edward Callaway | Osakada and Callaway (2013)<sup>59</sup> |
| HEK293T-TVA | Gift from Dr. Edward Callaway | Osakada and Callaway (2013)<sup>59</sup> |
| Experimental Models: Organisms/Strains |  |  |
| Mouse: Rbp4-Cre | MMRRC | MMRRC: 037128 |
| Mouse: Tnnt1-Cre | This paper | CDB0152E |
| Mouse: TIT2L-GC6s-ICL-tTA2 (Ai162) | The Jackson Laboratory | JAX: 031562 |
| Recombinant DNA |  |  |
| pAAV-CAG-FLEX-TCB | Miyamichi et al. (2013)<sup>47</sup> | Addgene_48332 |
| pAAV-CAG-FLEX-RG | Miyamichi et al. (2013)<sup>47</sup> | Addgene_48333 |
| Mouse whole brain cDNA | GenoStaff | Cat:MD01 |
| Oligonucleotides |  |  |
| Primers for RNA probes | This paper | N/A |
| Software and Algorithms |  |  |
| MATLAB | MathWorks | http://www.mathworks.com/<br>RRID: SCR_001622 |
| Photoshop | Adobe | N/A |
| Illustrator | Adobe | N/A |
| Inscopix Data Processing Software (IDPS) | Inscopix Inc. | N/A |
| Inscopix Data Acquisition Software (IDAS) | Inscopix Inc. | N/A |
| ilastik | Berg et al. (2019)<sup>60</sup> | https://www.ilastik.org/ |
| AMaSiNe | Song et al. (2020)<sup>48</sup> | https://github.com/cogilab/AMaSiNe |
| WaveSurfer | Howard Hughes Medical Institute | https://wavesurfer.janelia.org/ |
| MIN1PIPE | Lu et al. (2018)<sup>61</sup> | https://github.com/JinghaoLu/MIN1PIPE |
| Other |  |  |
| Inscopix nVista | Inscopix | RRID: SCR_017407 |
| Camera | Imaging source | DMK 33UX174 |

## METHOD DETAILS

### Animals

All experimental procedures were approved by the Institutional Animal Care and Use Committee of the RIKEN Kobe Branch. Animals were housed under a 12-h dark/light cycle with *ad libitum* access to food and water. Wild-type FVB mice were purchased from CLEA Japan, Inc. (Tokyo, Japan) for backcrossing. *Ai162* mice (*TIT2L-GC6s-ICL-tTA2*, Jax#031562) were purchased from The Jackson Laboratory. *Rbp4-Cre* mice were obtained from the Mutant Mouse Regional Resource Center. *Tnnt1-Cre* mice were generated as detailed below (background strain: C57BL/6). For all experiments, F1 hybrids of C57BL/6j and FVB strains were used, as they exhibit robust maternal behavioral acquisition.^27^ *Rbp4-Cre* heterozygous female mice (2–5 months old) were used for rabies tracing experiments. *Rbp4-Cre; Ai162* double-heterozygous female mice (2–5 months old) were used for *in vivo* microendoscopic imaging of the OFC. *Tnnt1-Cre; Ai162* double-heterozygous female mice (2–5 months old) were used for *in vivo* microendoscopic imaging of the thalamic nuclei. *Tnnt1-Cre* heterozygous female mice (2–5 months old) were used for anterograde tracing experiments. Because this study aimed to characterize neural representations in virgin female mice during maternal behavioral acquisition, only female mice were used.

### Generation of the *Tnnt1-Cre* line

The *Tnnt1-Cre* line (Accession No. CDB0152E; listed at https://large.riken.jp/distribution/mutant-list.html) was generated using CRISPR/Cas9-mediated knock-in in zygotes, as previously described.^62^ A donor vector consisting of *T2A-iCre* was inserted immediately upstream of the stop codon of *Tnnt1*. Guide RNA (gRNA) target sites were designed using CRISPRdirect^63^ to target upstream and downstream regions of the stop codon (Figure 2A). For microinjection, a mixture containing two CRISPR RNAs (crRNAs; 50 ng/μL), trans-activating crRNA (tracrRNA; 200 ng/μL), donor vector (10 ng/μL), and Cas9 protein (100 ng/μL) was injected into the pronucleus of a C57BL/6 one-cell stage zygote. Of 227 injected zygotes, 72 F_0_ founders were obtained, seven of which were iCre-positive as identified using PCR and sequencing. The line was established from a single male harboring the targeted sequence. Genotyping PCR was performed using the following primers: Tnnt1-iCre-F: 5′-GGTGGGTATTTGGTGGACTTCCTG, Tnnt1-iCre-R: 5′-GATGGCTCTCCCAGGCAGTATG, and iCre-F: 5′-TACCAAGCTGGTGGAGAGATGGATC, as shown in Figure 2B.

For sequence analysis, we designed the following primers: 3′-junc-R: 5′-AGGTCATGTCCTGGCAGTCTCAGTCC and 5′-junc-F: 5′-GCCGCTGGAAGGGCTCTGGCGAA. PCR products generated with Tnnt1-iCre-F and 3′ junc-R and with 5′ junc-F and Tnnt1-iCre-R were subcloned into pCR Blunt II TOPO vector (Zero Blunt TOPO PCR Cloning Kit, ThermoFisher) and sequenced using M13-Forward and M13-Reverse primers. The following RNAs were purchased from FASMAC (Atsugi, Japan): Tnnt1 crRNA (5′-CUG GAA GUG AGA CUG CCA GGg uuu uag agc uau gcu guu uug), PITCh 3 crRNA (5′-GCA UCG UAC GCG UAC GUG UUg uuu uag agc uau gcu guu uug), and tracrRNA (5′-AAA CAG CAU AGC AAG UUA AAA UAA GGC UAG UCC GUU AUC AAC UUG AAA AAG UGG CAC CGA GUC GGU GCU).

Germline transmission of the *Tnnt1-iCre* allele was confirmed by genotyping of F_1_ mice. Histochemical analysis using *in situ* hybridization (ISH) confirmed that expression was consistent with the endogenous Tnnt1 mRNA pattern (Figure 2).

### Viral preparations

The following AAV vectors were generated by UNC viral core using the corresponding plasmids as described previously.^47^ Titers were estimated using quantitative PCR and are reported as genome particles (gp) per milliliter:

AAV serotype 5 *CAG-FLEx-TVA-mCherry (TC^b^)* (9.3 × 10^12^ gp/mL)^47^
AAV serotype 8 *CAG-FLEx-RG* (2.8 × 10^12^ gp/mL)

The following AAV vectors were obtained from the Canadian Neurophotonics Platform Viral Vector Core facility and Addgene, produced using the corresponding plasmids (Addgene #154867, #177328, and #55637):

AAV serotype 8 *hSyn-fDIO-hM4Di-mCherry* (6.8 × 10^12^ gp/mL)^57^
AAV serotype 8 *hSyn-fDIO-mCherry* (8.9 × 10^12^ gp/mL)
AAV serotype retro *EF1a-FlpO* (5.5 × 10^12^ gp/mL)^58^

Rabies Δ*G-GFP*+EnvA was prepared using B7GG and BHK-EnvA cells (gifted from Ed Callaway) following established protocols.^59^ The EnvA-pseudotyped RVΔ*G-GFP*+EnvA titer was estimated at 1.0 × 10^9^ infectious particles/mL based on serial dilutions and infection of HEK293-TVA800 cells.

### Stereotaxic injection

Stereotaxic coordinates for each target brain region were determined from the Allen Brain Atlas.^64^ Mice were anesthetized with 65 mg/kg ketamine (Daiichi-Sankyo) and 13 mg/kg xylazine (Sigma-Aldrich) via intraperitoneal injection and head-fixed in stereotaxic equipment (Narishige). For rabies tracing experiments (Figure 1; Figure S1), 100 nL of a 1:1 mixture of AAV5 *CAG-FLEx-TC^b^* and AAV8 *CAG-FLEx-RG* was injected into the OFC (coordinates relative to bregma: anterior 2.5 mm, lateral 1.2 mm, and depth 1.9 mm from the brain surface) at 50 nL/min using UMP3 pump regulated by Micro-4 (World Precision Instruments). After 2 weeks, 200 nL of rabies Δ*G-GFP*+EnvA was injected. To minimize infection along the injection trajectory, rabies virus was injected using a tilted approach (coordinates relative to bregma: anterior 2.5 mm, lateral 0.5 mm, and depth 1.9 mm, with a 15° tilt from vertical). For anterograde tracing experiments (Figure 2F–H), 100 nL of AAV5 *CAG-FLEx-TC^b^* was injected into either the unilateral MD (coordinates relative to bregma: posterior 0.8 mm, lateral 0.5 mm, and depth 3.1 mm from the brain surface) or SMT (coordinates relative to bregma: anterior 0.8 mm, lateral 1.0 mm, and depth 4.0 mm from the brain surface, with a 15° tilt to minimize leakage into the MD). For pharmacogenetic experiments (Figure 5), 200 nL of AAV8 *hSyn-fDIO-hM4Di-mCherry* or AAV8 *hSyn-fDIO-mCherry* was bilaterally injected into the MD or SMT, followed by 100 nL of retro-AAV *EF1a-FlpO* into the bilateral OFC. After injections, the incision was sutured, and animals were placed on a heating pad to facilitate recovery from anesthesia before being returned to their home cages.

### Histology and histochemistry

Mice were given an overdose of isoflurane and transcardially perfused with PBS, followed by 4% paraformaldehyde (PFA) in PBS. Brain tissues were post-fixed in 4% PFA in PBS overnight at 4°C, cryoprotected with 30% sucrose solution in PBS at 4°C for 24–48 h, and embedded in O.C.T. compound (Tissue-Tek, cat#4583). For quantitative analysis of trans-synaptic tracing, 40-μm coronal sections of the whole brain were collected using cryostat (model #CM1860; Leica). Free-floating slices were incubated at room temperature with gentle agitation as follows: 2 h in blocking solution (5% heat-inactivated goat serum, 0.4% Triton X-100 in PBS); overnight in primary antibody diluted 1:1000 in blocking solution (mouse anti-GFP, GFP-1010, Aves Labs; or anti-RFP, 5f8, Chromotek); 2–3 h in secondary antibody diluted 1:500 (donkey anti-chicken-IgY Alexa488 or goat anti-rat IgG Cy3, Jackson ImmunoResearch); and 15 min in 2.5 μg/mL of DAPI (Santa Cruz, Cat #sc-3598) in PBS. Sections were mounted on slides and cover-slipped with mounting medium (Fluoromount; Diagnostic BioSystems).

Sections were imaged using Olympus BX53 microscope with a 4× (NA 0.16) or 10× (NA 0.4) objective lens equipped with cooled CCD camera (DP80; Olympus) or with Zeiss Axio Scan.Z1 using a 10× (NA 0.45) objective lens. Unless otherwise noted, every second (Figure 1; Figure S1) or third (Figure 2F–H; Figures S2 and S4) coronal brain section was analyzed for quantification.

### Rabies trans-synaptic tracing from the OFC

Animals were sacrificed 5 days after rabies injection (see **Stereotaxic injection**). For quantification, every second 40-μm coronal section of the whole brain was imaged using a 10× (NA 0.45) objective lens on Zeiss Axio Scan.Z1. Starter and input cells were detected semi-automatically using open source software, including ilastik^60^ and AMaSiNe,^48^ with custom-written MATLAB scripts.

Initially, ilastik was trained on a subset of slices to identify GFP and mCherry signals. Binary masks for GFP and mCherry channels were generated, and morphological opening in MATLAB was applied to remove noise. GFP+ cells were identified and converted into binary masks for detection in AMaSiNe. Starter cells were defined as overlapping GFP+ and mCherry+ signals, and only signals larger than five pixels were counted. Detected cells were reconstructed according to their spatial coordinates and annotated to brain regions using AMaSiNe. All automatic detections and regional assignments were manually inspected and corrected to ensure accuracy.

For analysis of long-range inputs, regions containing starter cells, such as the mOFC, M2, and PL, were excluded to avoid Cre-independent nonspecific labeling of rabies-GFP.^47^

### Axon tracing from the SMT/MD

AAV5 *CAG-FLEx-TC^b^* was injected into the SMT or MD (see **Stereotaxic injection**). Animals were euthanized more than 3 weeks after injection. To enhance axonal signals, sections were stained with anti-RFP (Chromotek). Every third 40-μm coronal section was imaged across the whole brain using 10× objective lens on Zeiss Axio Scan.Z1. Soma and axons were counted semi-automatically using ilastik and AMaSiNe, with custom-written MATLAB scripts. Briefly, ilastik was trained to distinguish axon fibers from soma, and binary masks for the mCherry channel were generated. Soma detection followed the same approach described for rabies tracing (Figure S2). For axon quantification, each pixel was treated as an individual “cell” in AMaSiNe and annotated to its corresponding brain regions to calculate axon density (Figure 2G, H). To account for individual variation in AAV transduction, axon density was normalized to the number of mCherry-positive cells per animal. In Figure 2H, the region with the highest mean axon density was designated as 1.

### Fluorescent ISH

Fluorescent ISH was performed as previously described.^65^ In brief, mice were anesthetized with sodium pentobarbital and perfused with PBS followed by 4% PFA in PBS. Brains were post-fixed in 4% PFA overnight. Thirty-micron coronal brain sections were cut using cryostat (Leica) and mounted on MAS-coated glass slides (Matsunami). To generate cRNA probes, DNA templates were amplified using PCR from the C57BL/6j mouse genome or whole-brain cDNA (Genostaff, cat#MD-01). A T3 RNA polymerase recognition site (5′-AATTAACCCTCACTAAAGGG) was added to the 3′ end of reverse primers. The primer sets for cRNA probe generation were as follows (forward primer first, reverse primer second):

*Tnnt1* 5′-AGCAGGCAGAAGATGAGGAA; 5′-TGGGGGCACTTTATTTTGAG
*vGluT2*-1 5′-TAGCTTCCTCTGTCCGTGGT; 5′-GGGCCAAAATCCTTTGTTTT
*vGluT2*-2 5′-CCACCAAATCTTACGGTGCT; 5′-GGAGCATACCCCTCCCTTTA
*vGluT2*-3 5′-CTCCCCCATTCACTACCTGA; 5′-GGTCAGGAGTGGTTTGCATT

DNA templates (500–1000 ng) amplified using PCR were subjected to *in vitro* transcription with DIG (cat#11277073910)-RNA labeling mix and T3 RNA polymerase (cat#11031163001) according to the manufacturer’s instructions (Roche Applied Science). For *vGluT2*, three independent RNA probes were mixed to increase the signal-to-noise ratio.

For ISH combined with anti-GFP staining, after hybridization and washing, sections were incubated overnight with horseradish peroxidase-conjugated anti-DIG (Roche Applied Science cat#11207733910, 1:500) and anti-GFP (Aves Labs cat#GFP-1010, 1:500) antibodies. Signals were amplified using TSA-plus Cyanine 3 (AKOYA Bioscience, NEL744001KT, 1:70 in 1× plus amplification diluent) for 25 min, followed by washing, and GFP-positive cells were visualized with anti-chicken Alexa Fluor 488 (Jackson Immuno Research cat#703-545-155, 1:250).

### Pup retrieval assay and co-housing

For microendoscopic recordings (Figures 3–5), animals were placed in their home cages (191 × 376 × 163 mm) with standard wood-chip bedding for at least 1 day before the assay. Before the first imaging session, animals underwent at least two habituation sessions, during which they were connected to the imaging rigs for 5–10 min. Retrieval assays were conducted once daily for three consecutive days. The assay began by introducing two pups (postnatal days 1–3 on experimental day 1) into opposite corners of the nest. If both pups were successfully retrieved, two additional pups were placed in the same corners. Each session lasted 6 min, and two imaging sessions were conducted per animal, separated by a minimum interval of 1 min. After the first day’s retrieval assay, the tested virgin female was cohoused with the mother and pups. On subsequent days, the mother and pups were temporarily removed during imaging, and a 5–10-min habituation period was included before the session. The same mother and pups were used across experimental days. The first session in which an animal retrieved more than 10 pups was defined as acquisition phase 1 (AP1), and the following day was defined as acquisition phase 2 (AP2).

### *In vivo* microendoscopic imaging

For microendoscopic recordings, ProView GRIN lens (500-μm diameter, 4 or 6.1 mm length; Inscopix) was implanted in target regions using the following stereotaxic coordinates: OFC, anterior −2.5 mm and lateral 1.2 mm relative to bregma, depth 1.8 mm from the brain surface; MD, anterior 0.8 mm and lateral 1 mm relative to bregma, depth 3.0 mm from the brain surface with a 10° tilt; SMT, anterior 0.8 mm and lateral 1.2 mm relative to bregma, depth 4.0 mm from the brain surface with a 15° tilt. Mice were anesthetized with 65 mg/kg ketamine (Daiichi-Sankyo) and 13 mg/kg xylazine (Sigma-Aldrich) via intraperitoneal injection and secured in stereotaxic equipment (Narishige). A 1-mm diameter craniotomy was made over the target area, and residual bone and dura were carefully removed using fine forceps. Brain tissue was aspirated to the depths of 1, 2, or 3 mm for the OFC, MD, or SMT, respectively. The GRIN lens, mounted on ProView lens holder and connected to Inscopix nVista, was slowly lowered into the brain while monitoring GCaMP6s fluorescence. Once GCaMP6s signals at the desired depth were confirmed, the lens was fixed with Super-Bond (Sun Medical) and the skull-lens interface was sealed. A metal head bar (Narishige, CF-10) was affixed to facilitate microscope attachment and detachment. After curing, the camera and lens holder were detached, and the exposed lens surface was protected with Kwik-Kast (WPI). After more than 3 weeks of recovery, mice were anesthetized, placed in the stereotaxic equipment, and the focal plane was adjusted to bring GCaMP6s-expressing cells into focus. A baseplate (Inscopix) was permanently affixed with Super-Bond. Following at least one additional week of recovery, mice were habituated to the microendoscope in their home cage for 5–10 min, repeated at least twice prior to imaging.

Imaging was performed using Inscopix nVista system (Inscopix). The focal plane was adjusted only at the start of each day, and imaging proceeded without further refocusing during sessions. Images (1080 × 1080 pixels) were acquired at 10 Hz using nVista HD software (Inscopix), with LED power set at 0.4–1 mW/mm^2^ and gain at 2.0–3.0. Timestamps for imaging frames and the camera were collected using WaveSurfer (https://wavesurfer.janelia.org/) for alignment. Two imaging sessions were conducted per day, each lasting 6 min, with short intervals between sessions. Imaging data were cropped to 800 × 700 pixels and exported as .tiff files using Inscopix Data Processing Software. Putative cell bodies were identified and neural signals extracted using MIN1PIPE v2 (https://github.com/JinghaoLu/MIN1PIPE^61^) with a spatial down-sampling rate of 2. All extracted traces were manually inspected, and cells with abnormal morphology or overlapping signals from adjacent cells were excluded. Relative changes in calcium fluorescence (*F*) were calculated as *ΔF/F_0_ = (F−F_0_)/F_0_* (where *F_0_* was defined as the median fluorescence of the entire trace). *ΔF/F* was normalized within each cell.

Behavioral videos were acquired at 20 Hz using a camera (Imaging source, DMK 33UX174). Transistor–transistor logic pulses generated by WaveSurfer were used to synchronize behavioral tracking with microendoscopic imaging.

### Pharmacogenetics

To silence neural activity via hM4Di (Figure 5), 100 μL of CNO (0.5 mg/mL, Sigma; C0832) was injected intraperitoneally after two 6-min imaging sessions (pre-CNO sessions). Two additional imaging sessions were performed 30 min after CNO administration (post-CNO sessions).

### Clustering analysis for imaging data of SMT^Tnnt^^1^ and MD^Tnnt^^1^

To assess functional heterogeneity, we performed clustering analysis on averaged responses during pup retrieval trials in SMT^Tnnt1^ or MD^Tnnt1^ neurons. Activity traces from −2 to 8 s relative to pup contact were trial-averaged and pooled across AP1 and AP2, yielding a 734 × 100 data matrix. Dimensionality reduction was performed using principal component analysis (PCA, MATLAB *pca*), and the first 11 principal components (PCs), explaining >95% of the variance, were retained. *K*-means clustering was then applied to the PC scores (five clusters, fixed random seed). To quantify response magnitude (Figures 3 and 4), response windows relative to pup contact were defined as follows: Cluster 1, −1 to +2 s; Clusters 2 and 4, 0 to 4 s; Cluster 3, 2.5 to 6 s. Baseline windows were defined as: Cluster 1, −3 to −2 s; Clusters 2–5, −3 to −1 s. Response amplitudes were quantified as the AUC during the response window, normalized to the mean baseline activity of each cluster.

### Decoder analysis

To test whether neural activity could predict behavioral events (pup contact, onset of retrieval, and completion) (Figures 3H and 4H), binary SVM classifiers were trained using a one-vs.-one multiclass strategy (MATLAB *fitcecoc* function). The first 10 retrieval trials were used, with the number of trials matched across conditions. For each iteration, data were split into training and test sets at a 4:1 ratio. Calcium signals were convolved within a 500-ms window centered on event onset. Training was performed using fivefold cross-validation, and predictions were aggregated by majority vote across classifiers. Decoding accuracy was defined as the percentage of correctly classified test samples. To obtain stable estimates, classification was repeated 5,000 times, with a random subset of ROIs selected for each iteration. Final accuracy was reported as the mean across iterations.

### Clustering analysis for OFC^Rbp^^4^ imaging data

To identify functional subtypes (clusters) in OFC^Rbp4^ neuron activity, clustering analysis was performed as described above. Activity traces from −2 to 8 s relative to pup contact were trial-averaged and pooled across three groups (Control, Gi-SMT, and Gi-MD) and two conditions (Pre- and Post-CNO), yielding a 1674 × 100 data matrix. PCA was applied using MATLAB *pca* function, and the first 9 PCs, explaining >95% of the variance, were retained. *K*-means clustering was then performed on the PC scores (six clusters, fixed random seed). To quantify response magnitude (Figure 5), response windows relative to pup contact were defined as follows: Cluster 1, −1 to 1 s; Clusters 2, 3, and 5, 0 to 4 s; Cluster 4, 0 to 6 s. Baseline windows were defined as follows: Cluster 1, −3 to −2 s; Clusters 2–5, −2 to −0.5 s. Response amplitudes were quantified as the AUC during the response window, normalized to the mean baseline activity of each cluster.

### Quantification and statistics

All statistical analyses were performed using custom MATLAB scripts. Tests were two-tailed, with significance defined as p < 0.05. Sample sizes and test types are indicated in the figure legends. Data are reported as mean ± standard error of the mean (SEM), unless otherwise indicated.

